# Localization of balanced chromosome translocation breakpoints by long-read sequencing on the Oxford Nanopore platform

**DOI:** 10.1101/419531

**Authors:** Liang Hu, Fan Liang, Dehua Cheng, Zhiyuan Zhang, Guoliang Yu, Jianjun Zha, Yang Wang, Feng Wang, Yueqiu Tan, Depeng Wang, Kai Wang, Ge Lin

## Abstract

Structural variants (SVs) in genomes, including translocations, inversions, insertions, deletions and duplications, remain difficult to be detected reliably by traditional genomic technologies. In particular, balanced translocations and inversions cannot be detected by microarrays since they do not alter chromosome copy numbers; they cannot be reliably detected by short-read sequencing either, since many breakpoints are located within repetitive regions of the genome that are unmappable by short reads. However, the detection and the precise localization of breakpoints at the nucleotide level are important to study the genetic causes in patients carrying balanced translocations or inversions. Long-read sequencing techniques, such as the Oxford Nanopore Technology (ONT), may detect these SVs in a more direct, efficient and accurate manner. In this study, we applied whole-genome long-read sequencing on the Oxford Nanopore GridION sequencer to detect the breakpoints from 6 carriers of balanced translocations and one carrier of inversion, where SVs had initially been detected by karyotyping at the chromosome level. The results showed that all the balanced translocations were detected with ∼10X coverage and were consistent with the karyotyping results. PCR and Sanger sequencing confirmed 8 of the 14 breakpoints to single base resolution, yet other breakpoints cannot be refined to single-base due to their localization at highly repetitive regions or pericentromeric regions, or due to the possible presence of local deletions/duplications. Our results indicate that low-coverage whole-genome sequencing is an ideal tool for the precise localization of most translocation breakpoints and may provide haplotype information on the breakpoint-linked SNPs, which may be widely applied in SV detection, therapeutic monitoring, assisted reproduction technology (ART) and preimplantation genetic diagnosis (PGD).

## Introduction

Structural variants (SVs), including translocations, inversions, deletions and duplications, account for a large number of variable bases, potentially leading to human genetic disorders due to disruption or dosage changes of functionally important genes[1-4]. In particular, balanced chromosome translocation, a common type of structural variants (SVs), is caused by the interchange of chromosomal segments between chromosomes, whereas inversions occurs when a single chromosome undergoes breakage and rearrangement within itself. In most cases, the altered karyotype has no immediately observable phenotype because an overall gene copy number is maintained, despite the possibility of the alterations of regulatory elements that influence gene expression. However, in a minority of cases, the breakpoints of translocation/inversion disrupt the gene structures, causing loss of function in genes associated with various diseases including infertility, disease syndromes, and congenital abnormalities[5-12]. Balanced translocation occurs in approximately 0.2% of the human population and 2.2% in patients who experience a history of recurrent miscarriages or repeated in vitro fertilization (IVF) failure[13, 14].

In somatic cells, balanced translocations can proceed through mitosis and replicate faithfully. However, during meiosis, chromosomes carrying balanced translocation are prone to abnormal segregation, leading to a variety of unbalanced translocation up to approximately 70%, which are derivatives with duplication and deletion of terminal sequences on either side of the breakpoint[15, 16]. Thus, parents who carry a balanced translocation in genome would face with a common reproductive outcome such as severe delay in successful conception, multiple miscarriages and occasionally children with a chromosome disease syndrome[17]. These couples commonly seek assisted reproductive technology (ART) and preimplantation genetic diagnosis (PGD) which aim to identify balanced euploid embryos for intrauterine transplantation and subsequently developing to a healthy infant[16, 18]. Hence, the precise location of translocation breakpoints is of great importance to increase the success rates of ART, considering the economic and psychological burdens to the families.

Karyotype analysis is a powerful, cost-effective, and long-established technology that remains widely applied in cytogenetics[19]. Although it has limited sensitivity and resolution, it can be a valuable diagnostic tool that provides input in genetic counseling for infertile patients[19, 20]. However, the low-resolution of this method restricted that it cannot identify cryptic balanced translocations and cannot identify the breakpoints precisely to infer the functional consequences of these chromosomal abnormalities.

So far, traditional methods to determine breakpoints of translocations include fluorescence *in situ* hybridization (FISH), Southern blot hybridization, inverse PCR and long-range PCR. These techniques are all time-consuming, expensive, difficult to provide information about the breakpoint-linked SNPs, and often fail to reach a diagnosis[21]. With the advances in sequencing technology, next-generation sequencing (NGS) have greatly expanded testing options, and provide a new avenue for translocation analysis and breakpoints detection[5, 21-23]. In addition, a “MicroSeq-PGD” method which combined chromosome micro-dissection and NGS can characterize the DNA sequence of the translocation breakpoints[24]. However, accurate detection of breakpoints using NGS has natural limitations due to the low mappability complex repetitive regions of the genome by the short reads (typically <150bp).

Nanopore sequencing, a single-molecule long-read sequencing technology, was first proposed by Deamer, Branton and Church, independently[25]. With the rapid improvements of nanopore sequencing technology and the development of bioinformatic tools designed for such data, it is becoming a valuable tool for clinical testing that addresses limitations from short-read sequencing. Though nanopore sequencing technology still has high error rate, which currently precludes their application in detecting single nucleotide substitutions and small frameshift mutations[26] under low coverage, the long read length (>10kb on average) enables greatly improved detection of SVs even in repetitive regions and provides an ideal tool for the detection of translocation breakpoints.

The long reads are especially useful in resolving breakpoints in repetitive regions of the genome with transposable elements. Transposable elements, including DNA transposons and retrotransposons, are major contributors to genomic instability. Endogenous retroviruses, long interspersed elements (LINEs), and short interspersed elements (SINEs) belong to retrotransposon. Alu element, one of the SINEs, is the most successful retrotransposon in primate genomes, composing 10% of the human genome[27]. Genomic rearrangements induced by Alu insertion account for approximately 0.1% of human diseases and genomic deletions by Alu recombination-mediated deletions (ARMD) are responsible for approximately 0.3% of human genetic disorders[28-30].

The long reads are also useful to resolve haplotypes between a translocation and the nearby SNPs or indels, which is of special importance in preimplantation genetic diagnosis (PGD). Due to the presence of allelic drop-out when assaying single cells in PGD, the markers along a very long stretch of DNA can indicate whether the chromosome carries translocation or not in each embryo. This method, preimplantation genetic haplotyping (PGH), is a simple, efficient, and widely used method to identify and distinguish between all forms of the translocation status in cleavage stage embryos prior to implantation[31]. Generally speaking, haplotypes are established using informative polymorphic markers which covered ±2Mb around the breakpoints. Meanwhile, these SNPs should be homozygous in the carrier’s parents or other family members.

In this study, we demonstrated the ability of Oxford Nanopore sequencing to detect translocations and refine their breakpoints, which were initially detected by conventional karyotyping. Fourteen breakpoints from seven carriers were detected successfully and most of them were mapped to single base resolution by Sanger sequencing. Meanwhile, we also obtained the haplotype information surrounding the breakpoint regions, which facilitates single-cell sequencing in preimplantation genetic diagnosis (PGD). Our results indicate that low-coverage whole-genome sequencing is an ideal tool for the precise localization of translocation breakpoints, which may be widely applied in SV detection, therapeutic monitoring, assisted reproduction technology (ART) and preimplantation genetic diagnosis (PGD).

## Material and Methods

### Samples

The study was approved by the Institutional Review Board of the CITIC-Xiangya Reproductive and Genetics Hospital, and written informed consent were obtained from all participants. A total of 7 patients, including 3 with long-standing infertility, were recruited at the CITIC-Xiangya Reproductive and Genetics Hospital. Among them, 6 balance translocations and 1 inversion were previously identified by karyotyping. The mean maternal age was 30.4 years (21–34 years), indicating a moderate risk of incidental aneuploidies. There are 3 female carriers and 4 male carriers. DNA was extracted using FineMag Blood DNA Kit (GENFINE BIOTECH) according to the manufacturer’s instructions.

### Library preparation and sequencing

5 μg genomic DNA was sheared to ∼5-25kb fragments using Megaruptor^®^ 2 (Diagenode, B06010002), size selected (10-30kb) with a Blue Pippin (Sage Science,MA) to ensure the removal of small DNA fragments. Subsequently, genomic libraries were prepared using the Ligation sequencing 1D kit (SQK-LSK108, Oxford Nanopore, UK). End-repair and dA-tailing of DNA fragments according to protocol recommendations was performed using the Ultra II End Prep module (NEB, E7546L). At last, the purified dA tailed sample, blunt/TA ligase master mix (#M0367, NEB), tethered 1D adapter mix using SQK-LSK108 (Oxford Nanopore Technologies) (ONT) were incubated and purified. Library was sequenced on R9.4 flowcells using GridION X5.

### SVs analysis

The raw sequencing output as FAST5 files were converted to FASTQ format using the MINKNOW local basecaller. SVs were called using a pipeline that combines NGMLR-sniffles and LAST-NanoSV. Briefly, long reads were aligned to human reference genome (hg19) by using NGMLR [32] (version 0.2.6) with ‘-x ont’ argument and LAST (version 912) separately, then SV call sets were performed by sniffles(1.0.6) with ‘report BND ignoresd q 0 genotype -n 10 -t 20 -l 50 -s 1’ and NanoSV [33] with ‘-c 1’ arguments. In order to improve sensitivity of translocation calling, a custom python scripts was developed to obtain all the split reads that were mapped to different chromosomes. Also, the alignment information about identity, mapping quality, matched place and matched length is retained. IGV[34] and Ribbon[35] were used for visual examination of translocations in target region. Inversions were detected by combining results of sniffles and NanoSV.

### Breakpoint verification

We designed PCR primers to detect the translocation breakpoints for each sample. Primer3-Plus (http://primer3plus.com/) was used for primer design. All primers used in this study were provided in Supplementary Table S1. PCR was performed using 2X Taq Plus Master Mix polymerase (P211-01/02/03, Vazyme), and the products were electrophoresed through a 1.0% agarose gel and sequenced by Sanger sequencing on an ABI3730XL sequencer (Applied Biosystems). PCR conditions are available on request.

### CNVs analysis

CNV analysis was performed by Xcavator, a software package for CNV identification from short and long reads of whole genome sequencing experiments[36]. For each sample, CNV was called by using the other six individuals as controls. During the sequencing process, as each read was randomly and independently sequenced from any location of the genome, the copy number of any genomic region could be estimated by counting the number of reads (read count) aligned to consecutive and non-overlapping windows of the genome. As a result of low sequencing coverages (≤ 10x), we used 1kb window size.

### Haplotype analysis

MarginPhase is a method that uses a Hidden Markov Model to partition long reads into haplotypes[37]. After we obtain candidate SVs by the combined pipeline described above, we get the ±2Mb sequences around the breakpoint. To identify mutations, SNP/indels was first called using SAMtools mpileup and bcftools. Finally, we generate haplotype calls using MarginPhase.

## Results

### Chromosomal analysis of carries with balanced translocations or inversion

We recruited 7 carriers of translocation in total in the study from CITIC-Xiangya Reproductive and Genetics Hospital (Table 1). These subjects were affected with either long-standing infertility, or had a history of recurrent miscarriage or had children bearing chromosome syndromes. About 5 ml blood from each carrier was extracted, and 2 ml was mixed with peripheral blood culture medium and cultured in an incubator at 37 °C. After 72 hours, harvested chromosome specimens were prepared and subject to a G-banding karyotype analysis by standard protocols, according to the International System for Human Cytogenetic Nomenclature. The results revealed that six of the carriers had reciprocal balanced translocations and the last one had an inversion translocation (Fig S1). We decided to perform whole-genome long-read sequencing on all subjects, to map the exact breakpoints. Based on the karyotyping results, we chose different analytical strategies and software tools to analyze the translocation breakpoints in the next step.

**Table 1.** The list of subjects analyzed in the current study and the details on the inferred breakpoints.

| Sample | Karyotype | Depth (X) | No. of mapped sequencing reads | No. of mapped sequencing bases | Coverage rate (%) | No. of spanning breakpoints reads | Breakpoint position (GRCh37) | Disrupted gene (breakpoint) |
| --- | --- | --- | --- | --- | --- | --- | --- | --- |
| DM17A2236 | 46,XY,t(6;8)(q25;q22) | 11.32 | 2,262,314 | 32,111,789,470 | 91.85 | 11 | 6:167281717<br>8:113696089 | Intergenic region<br>*CSMD3* |
| DM17A2237 | 46,XX,t(18;21)(q11;q11) | 10.31 | 2,316,017 | 29,746,593,714 | 93.44 | 11 | 18:28685658<br>21:29073597 | *DSCAS*<br>Intergenic region |
| DM17A2246 | 46,XX,t(8;22)(q24;q11) | 9.87 | 1,931,784 | 28,742,307,402 | 94.34 | 6 | 8:125495366<br>22:20326956~20327048 | *RNF139*<br>Intergenic region |
| DM17A2247 | 46,XY,t(11;22)(q23;q11) | 9.98 | 2,024,838 | 29,361,507,192 | 95.30 | 5 | 11:116683166<br>22:20326993 | Intergenic region<br>Intergenic region |
| DM17A2248 | 46,XX,inv(11)(q11q21) | 10.94 | 2,498,061 | 32,758,847,457 | 96.96 | 10 | 11:58265643<br>11:100448937 | Intergenic region<br>Intergenic region |
| DM17A2249 | 46,XY,t(2;18)(p13;q23) | 10.26 | 1,790,385 | 29,628,601,968 | 93.52 | 11 | 2:80320441<br>18:66637011 | *CTNNA2*<br>*CCDC102B* |
| DM17A2250 | 46,XX,t(3;9)(p13;p13) | 13.54 | 3,150,533 | 39,494,541,253 | 94.43 | 7 | 3:90504855<br>9:44216447 | centromere region<br>Intergenic region |

### DNA extraction and sequencing by GridION X5

For all subjects, genomic DNA was sheared to 10-20 kb fragments and DNA libraries were prepared and sequenced using standard protocols on the Oxford Nanopore GridION X5 sequencer. For all samples, mean identify and median identify of reads to the reference genome were mostly higher than 85% (Fig 1A). We obtained a total read bases of 32-44 Gb in each sample, with a mean length of 12.3-16.3 Kb and a depth of 9.87-13.54X (Fig 1B). These results suggested that we obtained high-quality sequencing data to facilitate downstream analysis. After sequencing, all the reads generated from each sample were aligned to the human reference genome (hg19), and used for subsequent downstream data analysis. The detailed results were summarized in Table S2.

**Figure 1.**
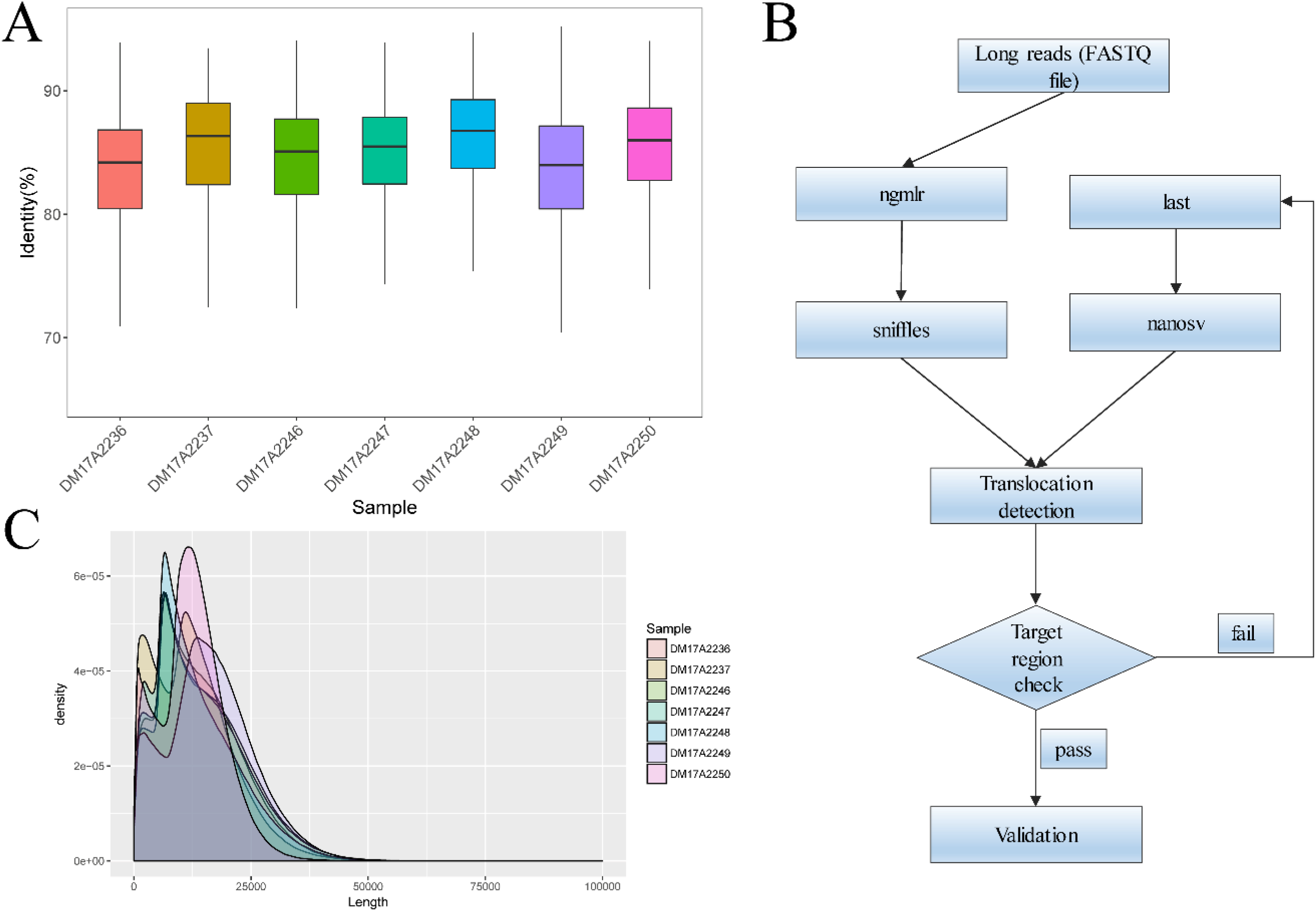
Quality control of the Oxford Nanopore long-read sequencing data. (A) Median identity of sequencing data to the reference genome is around 85% for all samples. (B) The mean length was 12.3-16.3 kb and the read N50 was 15.3-20.5kb for all samples. (C) The overall strategy for breakpoint analysis.

### Translocation detection and breakpoint characterization

We analyzed the long-read sequencing data obtained from Oxford Nanopore to detect the breakpoints in six individuals with balanced translocations and one individual with inversion using a custom bioinformatics pipeline that incorporate several existing tools (Fig 1C). This bioinformatics pipeline identified the potential breakpoints from the alignment data. For instance, 10 reads from sample DM17A2237 were used to locate the breakpoint to the point of chr18:28685658, whereas another 10 reads located the other breakpoint at position chr21:29073597 (Fig 2A). Through these long reads, we can accurately locate the breakpoints of this carrier at these two positions. Then, we designed PCR primers to verify the breakpoints by Sanger sequencing. We found that the translocation results were consistent with the karyotyping results (Fig 2B). All of the detailed breakpoints information and sequencing quality data from the 7 samples of carriers were summarized in Fig S2, Fig S3 and Table S2, respectively.

**Figure 2.**
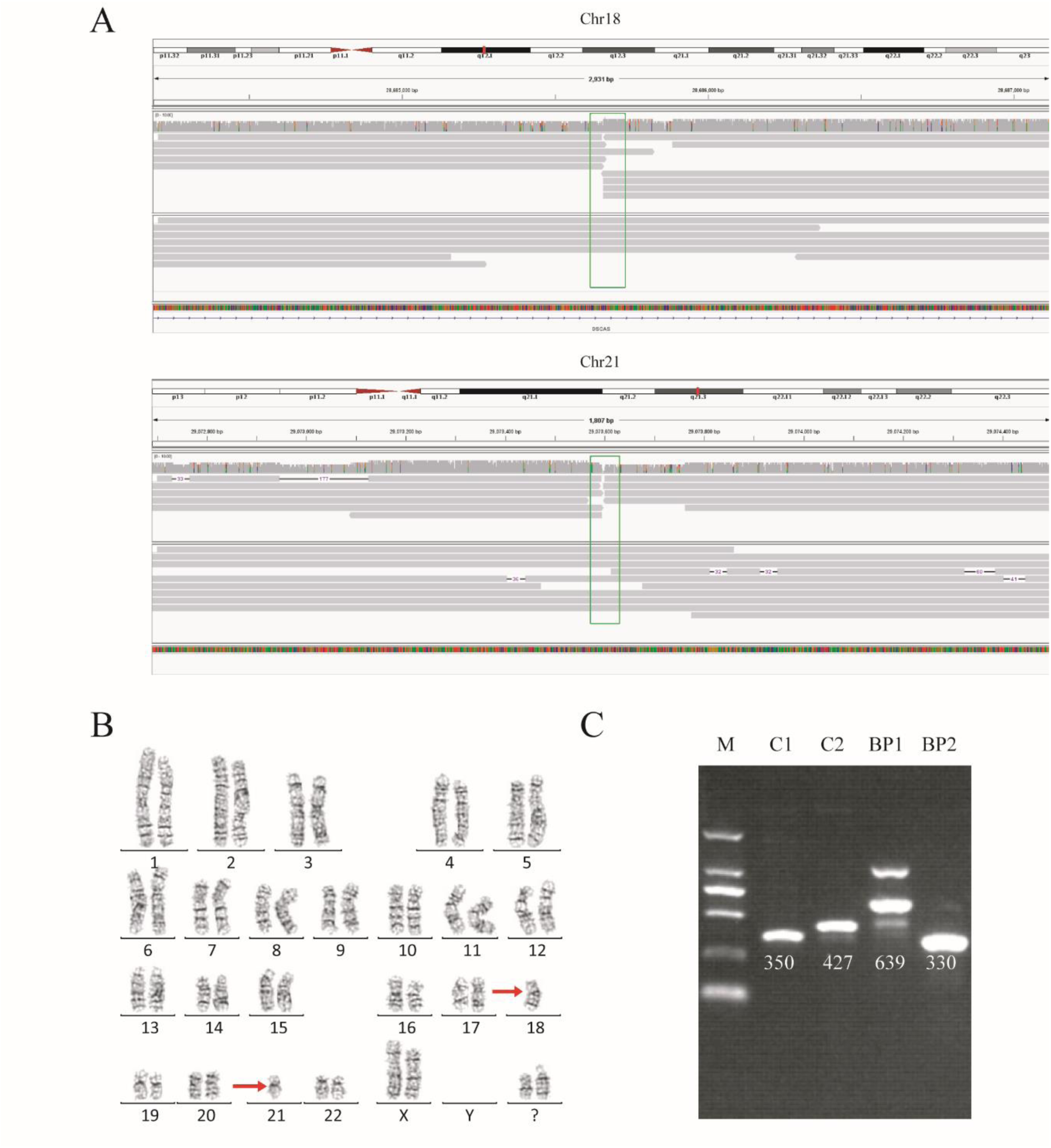
Balanced translocation by sequencing and karyotyping in subject DM17A2237. (A) Read mapping of the breakpoints for the balance translocation. DNA fragments were compared to human genome reference GRCh37/hg19, and the breakpoints were showed in Integrative Genomics Viewer (IGV). A total of 20 reads adjacent to the breakpoint were found. (B) Karyotype of carrier M17A2237. Karyotype analysis was determined from G-banding analysis by standard protocol. The karyotype result showed a rough region where the breakpoint occurred. (C) Polymerase chain reaction (PCR) analysis and Sanger sequencing to validate the breakpoints. Agarose-ethidium bromide gel showing the presence of two new bands created by rearrangement of chromosomal segments at breakpoints (BP1 and BP2). M=Marker, C=Control, BP=breakpoint. Primer information is available in Table S1.

Checking these breakpoints in the UCSC Genome Browser, we found that in sample DM17A2236, DM17A2246, DM17A2247 and DM17A2249, breakpoints were located within the introns of genes *CSMD3, AK129567, AK302545, RNF139* and *CCDC102B,* respectively. Therefore, these breakpoints disrupted the gene structures, causing exchange of materials between chromosomes, which impair gene function since a portion of the gene structure in one chromosome is moved to the other chromosome. Examination of medical records showed that altered karyotype had no obvious phenotypic consequences in the early years of the carrier, but almost all the carriers had a phenotype of primary infertility causing by failure of meiosis once they reach adulthood. Additionally, we also compared between two long-read alignment methods, and found that LAST could map more sequences than NGMLR, while NGMLR were able to map large gaps within long reads more reliably. In detection of translocation breakpoints, LAST had higher sensitivity but cost longer time. Considering these issues, we used NGMLR as the primary breakpoints detecting approach, and LAST as a supplementary approach to ensure more accurate results (Table S3). We also found that the aligned sequence of DM17A2246 was located at 22q11.21 with a 79 bp deletion (chr22:20656022-20656100). DM17A2247 had a gap of 33Kb (chr22:206326985-20656120). Furthermore, there are clusters of low-copy repeats (LCRs) in 22q11.21, which indicates that balanced translocation may occur preferentially at the site of LCRs cluster.

The genomic rearrangements caused by *Alu* elements could lead to genetic disorders such as hereditary disease, blood disorder, and neurological disorder[38]. Major *Alu* lineages are *AluJ, AluS*, and *AluY* are distinguishable from each other with 18 diagnostic nucleotides on their sequences[39]. In our study, we found that in sample DM17A2237, the breakpoint of chr18:28685658 occurred at *AluY* element; yet, in sample DM17A2250, the breakpoint of chr9:44216447 occurred at *AluSx3* element. Although these breakpoints did not compromise the structures of any genes, they may still be associated with infertility in these patients.

Interestingly, the subject DM17A2250 with a karyotype of 46,XX,t(3;9)(p13;p13) carries a balanced reciprocal translocation, which locates to chr3:90,490,057-90,504,855 and chr9:44,225,822, respectively. The breakpoint on chromosome 3 is very close to the acrocentric centromere. All the long reads show a clear breakpoint at chr3:90,504,854 consistent with the result of karyotyping, but it is not a typical Robertsonian translocation. Since most of translocations involving in acrocentric centromere are Robertsonian translocation, to the best of our knowledge, this is among the first report of t(3;9) that is not a usual Robertsonian translocation and has been mapped to single-base resolution by our approach.

### Inversion detection and breakpoint characterization

Similar to balanced translocations, inversion does not change chromosome copy number, and is difficult to detect by conventional short-read sequencing platforms, despite their functional consequences in medical genetics [40]. Here we successfully detected an inversion occurred in carrier DM17A2248 at chr11:58,255,398-58,293,470 and chr11:100,430,372-100,461,378 (Fig S2). After verification by PCR and Sanger sequencing, the breakpoints were finally mapped to chr11:58,265,643 and chr11:100,448,937, respectively, consistent with the karyotyping result. Our results demonstrated an example where long-read sequencing is capable of resolving complex breakpoints for inversions accurately.

### Breakpoint validation by Sanger sequencing

To further validate the exact translocation breakpoints and adjacent SNPs around the breakpoints, PCR reactions and Sanger sequencing were performed to map the breakpoint sequences at the level of individual bases. For translocations, we successfully identified the breakpoints in sample DM17A2236, DM17A2237, DM17A2248 and DM17A2249 by Sanger sequencing, but failed in DM17A2246, DM17A2247 and DM17A2250 (Fig S4). Because the approximate breakpoints of DM17A2246 and DM17A2247 are located in highly repetitive regions and the breakpoint of DM17A2250 is near a centromere, it is challenging to obtain a PCR product for these breakpoints after multiple failed attempts. Nevertheless, it is worth nothing that in sample DM17A2247, we have successfully obtained the target PCR bands from the normal chromosome without translocations, but no band was found based on the rearranged chromosomes (Fig S5), which reveals that there may be a deletion or larger insertion near the breakpoints to disturb the binding sites of our designed primers. The results above suggest the power of long-read sequencing in detecting precise locations of translocation breakpoints, while karyotype analysis can only provide rough results in the range of megabases. Therefore, long-read sequencing may be a more precise tool to detect translocation breakpoints that may complement or validate karyotyping results in clinical diagnosis settings.

### Haplotype detection

Haplotype identification of chromosome is of great importance to preimplantation genetic diagnosis (PGD), so that we can use adjacent SNP information to predict the presence or absence of balanced translocations in single-cell assays. Here we performed haplotype analysis by using the breakpoints as precise markers. Through these markers, we successfully found the informative SNPs near the breakpoint regions, to differentiate the parts of chromosomes involved in the translocation and the corresponding normal homologous chromosomes in sample DM17A2237 at a low-level coverage (10×) (Fig 3). The haplotypes will help to distinguish between embryos with balanced translocation and structurally normal chromosomes through PGD analysis, when the spouse of the carrier has normal karyotype. These results above demonstrate that it is possible to detect haplotype by low-coverage long-read sequencing, and obviously, more accurate haplotype information can be obtained by increasing sequence coverage.

**Figure 3.**
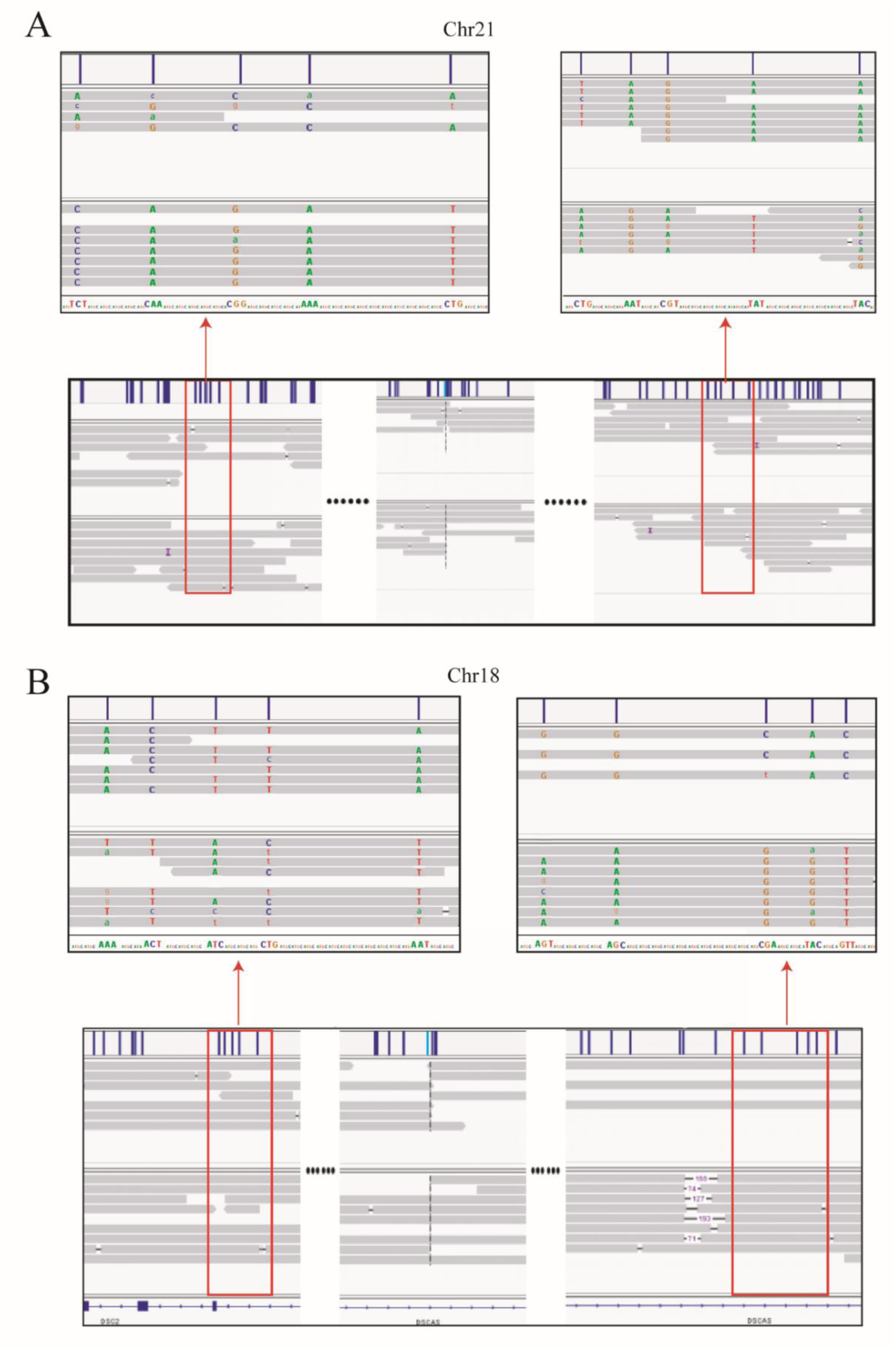
Long reads detected the haplotype around translocation breakpoints in sample DM17A2237. Using the breakpoints as anchoring markers, we obtained the ±2Mb sequences around the breakpoints. Through SNP calling and the MarginPhase tool, we phased the haplotypes around the breakpoints in chr18 and chr21. Majuscule letters represent accurate base information, while lower case letters represent fuzzy base information.

### Exploratory analysis of CNVs by low-coverage long-read sequencing

Copy number variant (CNV) is an important type of structural variants, and the identification of CNVs is also useful for clinical diagnose. In our exploratory analysis, CNVs from each subject were analyzed by using the sequencing data of the other six carriers as controls. The effective region (without gap) of reference genomic sequence was divided into blocks with a length of 100kb. Depth of each block was calculated and then region with a Z-score greater than 3 or less than -3 was defined as a potential CNV. Using this approach, we found that there were approximately 200 CNVs beyond 100Kb can be detected in each sample. Due to the relatively low whole-genome coverage, the minimum sequencing depth of coverage to detect CNVs is set as 2×. Additional simulation shows that a higher depth yields a better resolution, where more CNVs would be identified (Fig S6). Since our study focused on translocations that were already identified by karyotyping, we did not perform more detailed analysis on the CNVs. However, these results and simulations demonstrate that even with low-coverage data, long-read sequencing still has the ability to detect a large number of potential CNVs, and may be used to validate candidate CNVs that are detected from other platforms (such as SNP arrays).

## Discussion

Currently, the most widely used technology for clinical diagnose of chromosome translocation is karyotype analysis[41]. Although next-generation sequencing technology and gene chip technology offer high resolution, high sensitivity, high throughput [21, 23], those methods are not suitable for breakpoints detection of balanced translocation or inversions due to technical limitations. On the other hand, karyotype analysis is of low-resolution, yet the exact identification of breakpoints are often required to better understand how the translocations impact genes and phenotypes.

In this study, we used Oxford Nanopore Technology to analyze the genomic variations of 7 patients who suffer from long-standing reproductive disorder. All of the 7 patients carry chromosomal translocations in their genomes, among whom 6 have reciprocal balanced translocations and the last one has an inversion translocation. We have successfully identified and sequenced every breakpoint from the samples of the seven carriers by long-read sequencing. Among them, 8 breakpoints (4 carriers) were easily verified by Sanger sequencing (57.1%), while the other 6 breakpoints (3 carriers) failed in PCR amplification for Sanger sequencing (42.9%), because of the repetitive or centromere regions in the target sites. Nevertheless, all of these 14 breakpoints identified by long-read sequencing were consistent with their corresponding karyotype results. This finding provide strong evidence that long-read sequencing shows flexibility in sequence preference even if the breakpoints appeared in highly repetitive and complicated regions.

*Alu* repeats represent the largest family of mobile elements in the human genome, and continue to generate genomic diversity in several ways[42]. In our results, we found two breakpoints that occurred at *Alu* elements. Meanwhile, in sample DM17A2249, there is also a breakpoint in L1PA4, which is an repeat element. Repetitive elements have been implicated as the sites of chromosome instability, so our results suggest that these elements may be susceptible to generate balanced chromosomal translocation.

Robertsonian translocation formed by abnormal breakage and joining of two acrocentric chromosomes has an estimated 0.1% incidence rate in the general population[25]. Balanced non-RT, involving acrocentric centromere, is a rare event and only a few cases are reported. In our research, we first report a non-RT at t(3;9) and locate the breakpoints successfully. However, additional work are needed to complete sequencing of human genome, because the majority of sequence information near the acrocentric centromere is still unknown.

In addition, PCR identification of sample DM17A2249 and DM17A2248 showed clear target bands of the wild type copies at the breakpoints sites, but failed to get any band at least at one or both breakpoints in the homologous chromosomes carrying translocation. Reciprocal chromosome translocations are often accompanied by some additional rearrangements, such as deletions and duplications, involving only a few base pairs to megabases in extent. As previously reported, almost 50% balanced translocations show large deletions and duplications at the breakpoint junction[43, 44]. The failures of breakpoints identification by PCR in sample DM17A2249 and DM17A2248 may be due to the existence of this kind of rearrangements, for which a deletion leads to a loss of binding site by PCR primers or a large insertion makes the PCR product too long to obtain.

In conclusion, taking advantage of the long reads, low-coverage whole-genome sequencing could be a more efficient and powerful tool to analyze chromosomal translocations, compared with traditional methods such as FISH and NGS. Based on comparison to the karyotyping results and our Sanger sequencing results, we confirmed that nanopore sequencing exhibits high resolution and accuracy. We believe that long-read sequencing may play a more important role in chromosomal translocation analysis and breakpoints detection in the future, and offer valuable insights to assist the genetic diagnosis of reproduction and preimplantation.

## Acknowledgements

The authors want to thank patients who participated in this study to evaluate novel genomic approaches for improved genetic diagnosis of balanced translocations and inversions. We also thank the genetic counselors and clinical geneticists who interviewed the patients and collected DNA samples. This study was partially supported by National Key R&D Program of China (SQ2018YFC100084) and Merck Serono China Research Fund for Fertility Experts.

## Competing Interests

L.H., D.C., Y.T. and G.L. are employees of Reproductive & Genetic Hospital of CITIC–Xiangya. Z.Z., F.L., Y.W. and D.W. are employees and F.W. and K.W. are consultants of Grandomics Biosciences.

## Supplementary materials

**Figure S1.**
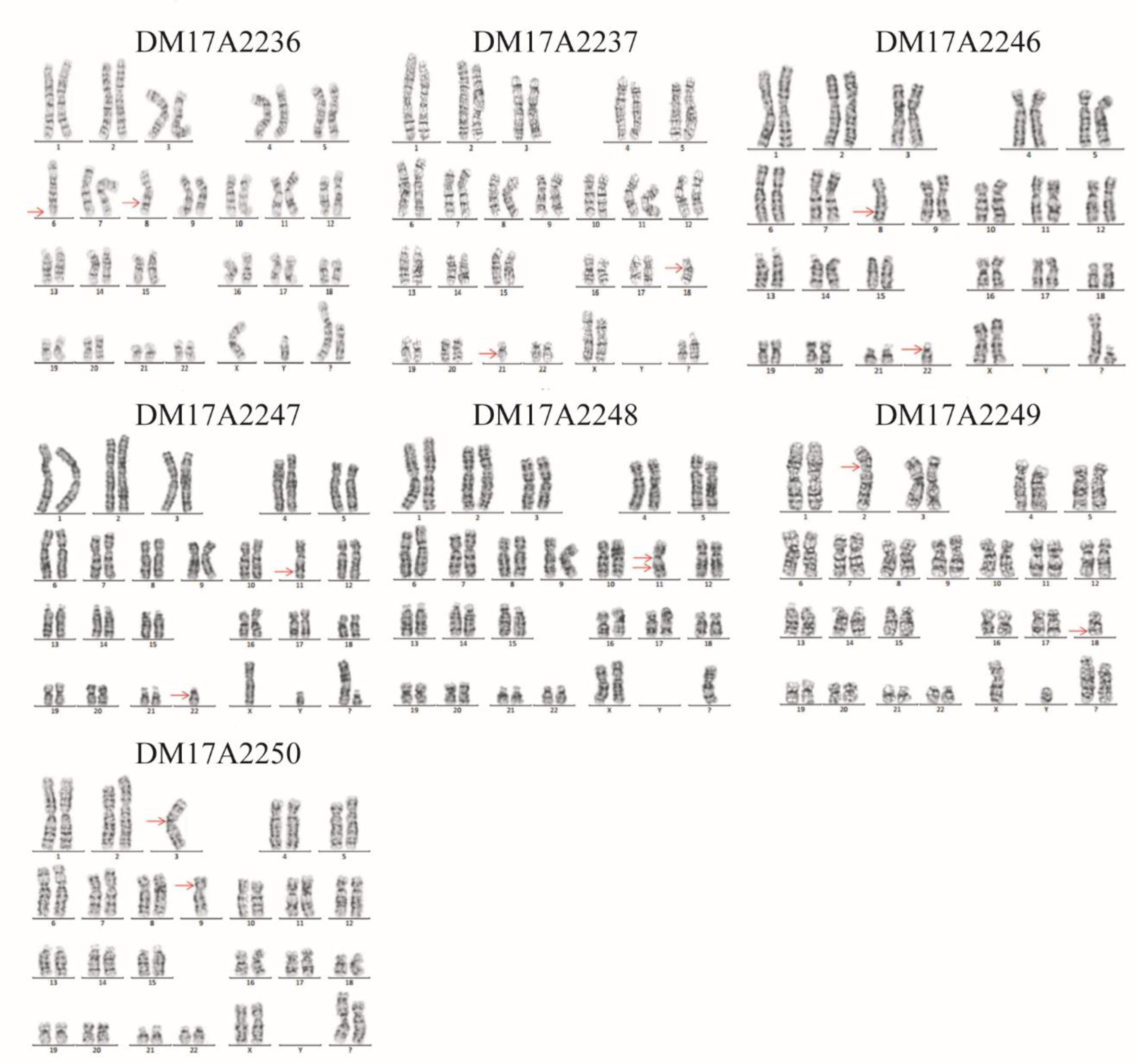
Karyotypes of 7 subjects. See Table 1 for details.

**Figure S2.**
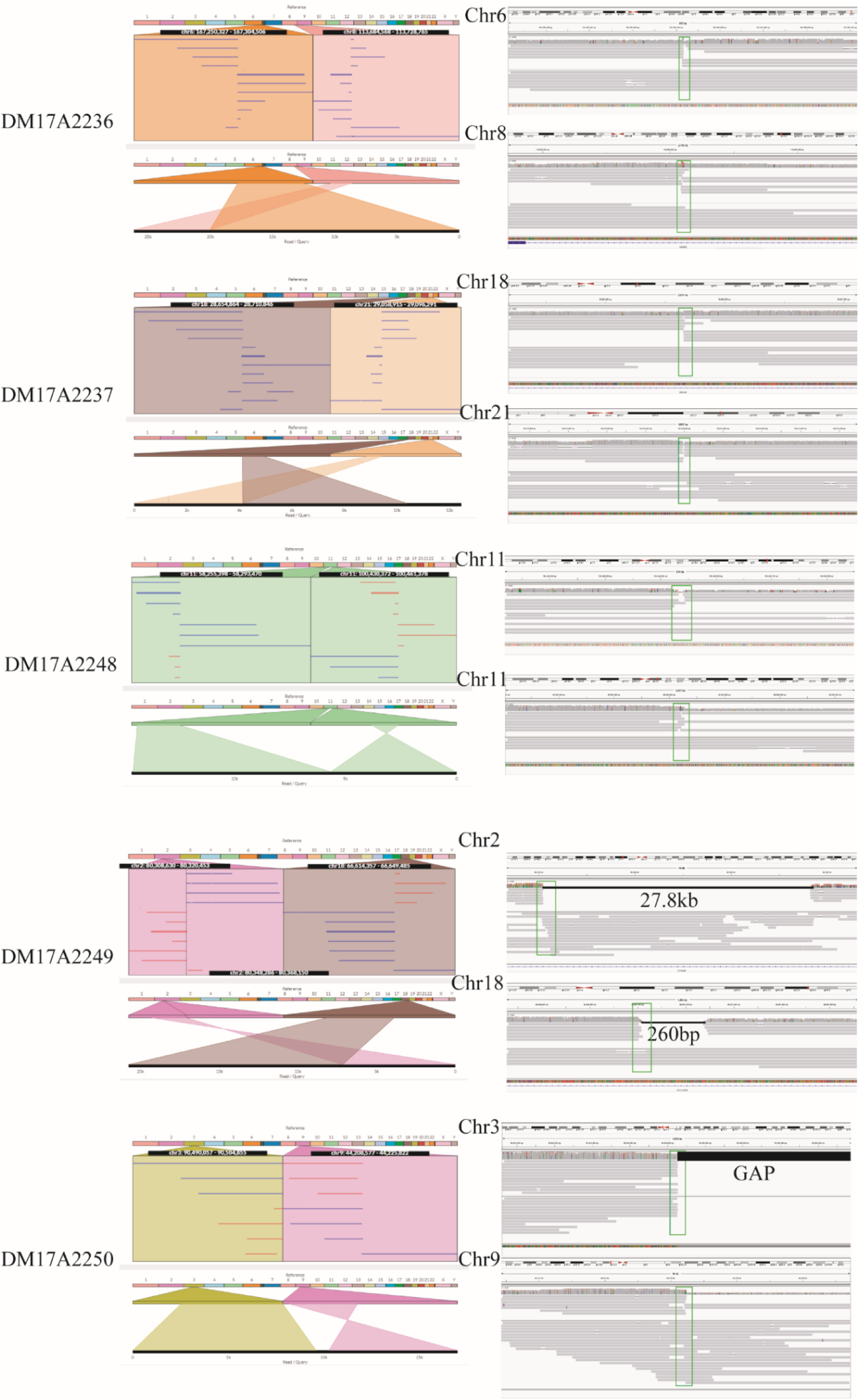
Translocation detection and analysis by long-read sequencing, as illustrated by Ribbon and IGV.

**Figure S3.**
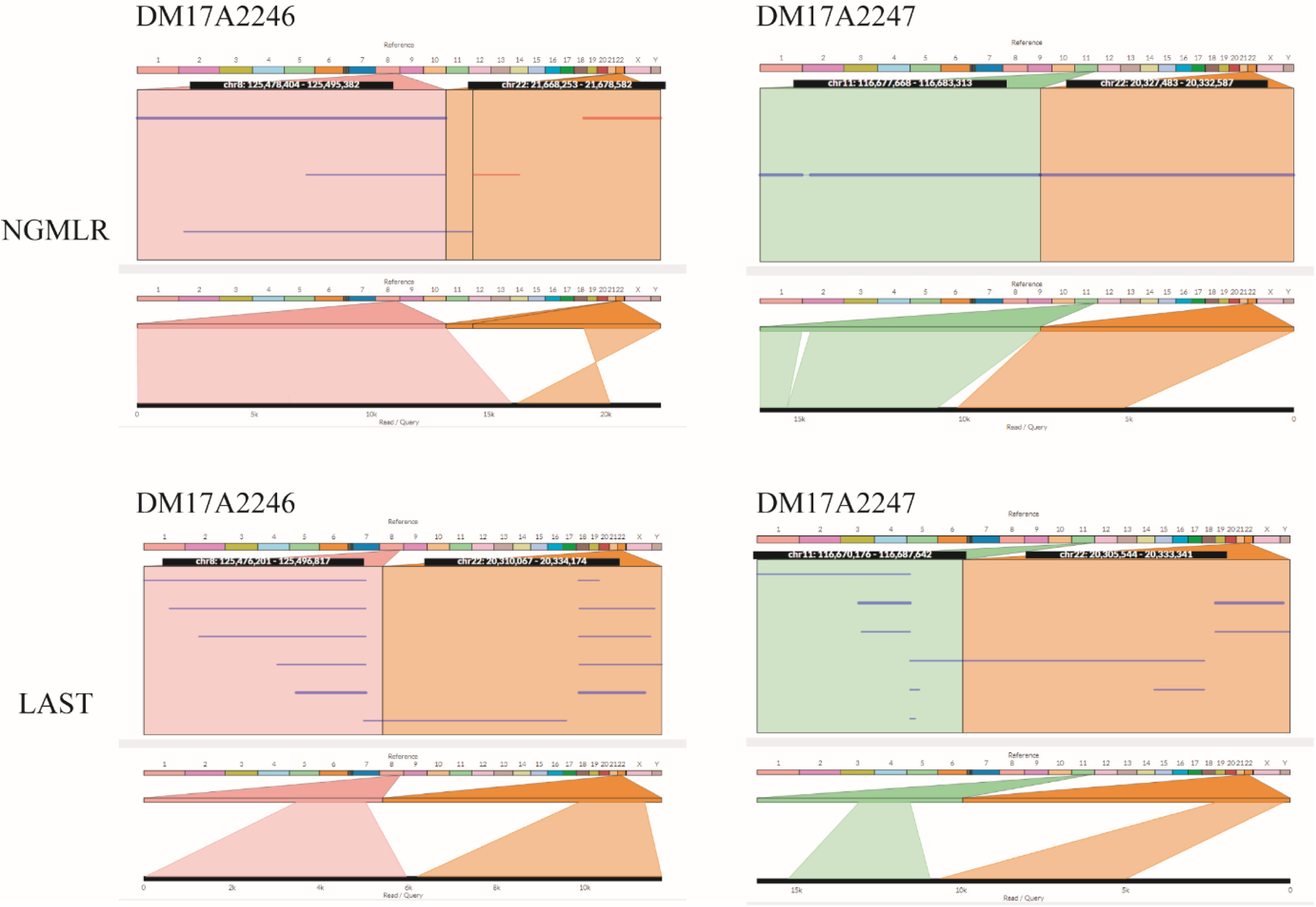
Translocation analysis by NGMLR and Last in DM17A2246 and DM17A2247, as illustrated by Ribbon.

**Figure S4.**
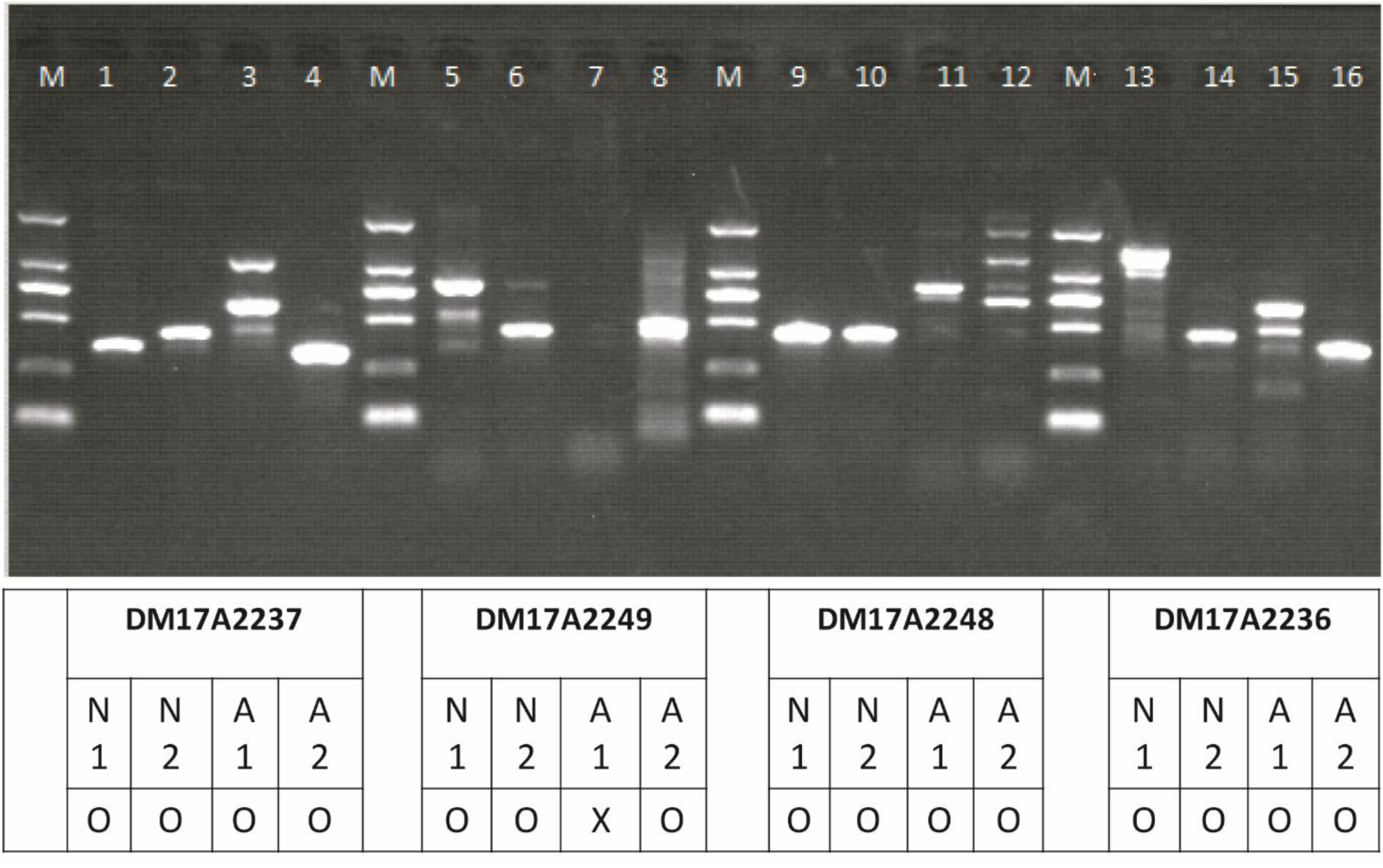
Verification of translocation breakpoints by PCR and Sanger sequencing.

**Figure S5.**
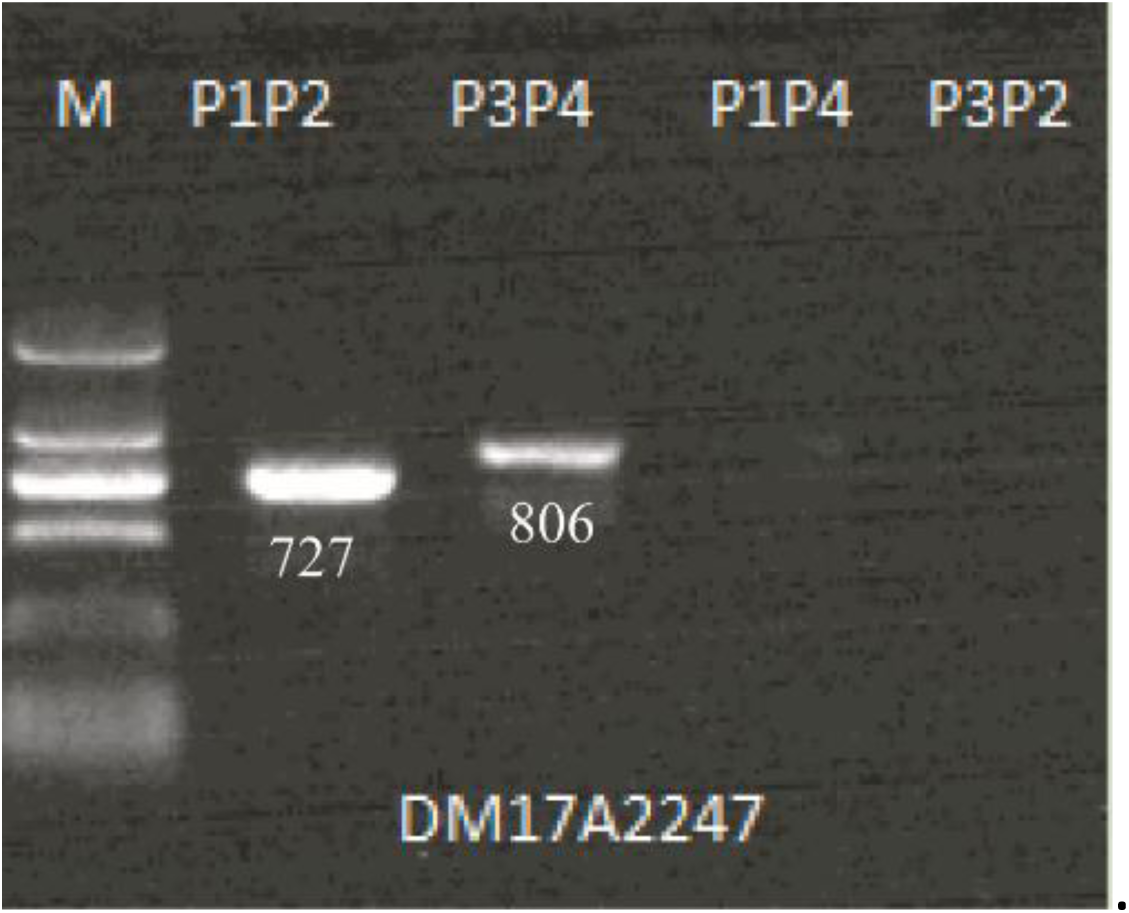
Verification of translocation breakpoints by PCR and Sanger sequencing in sample DM17A2247

**Figure S6.**
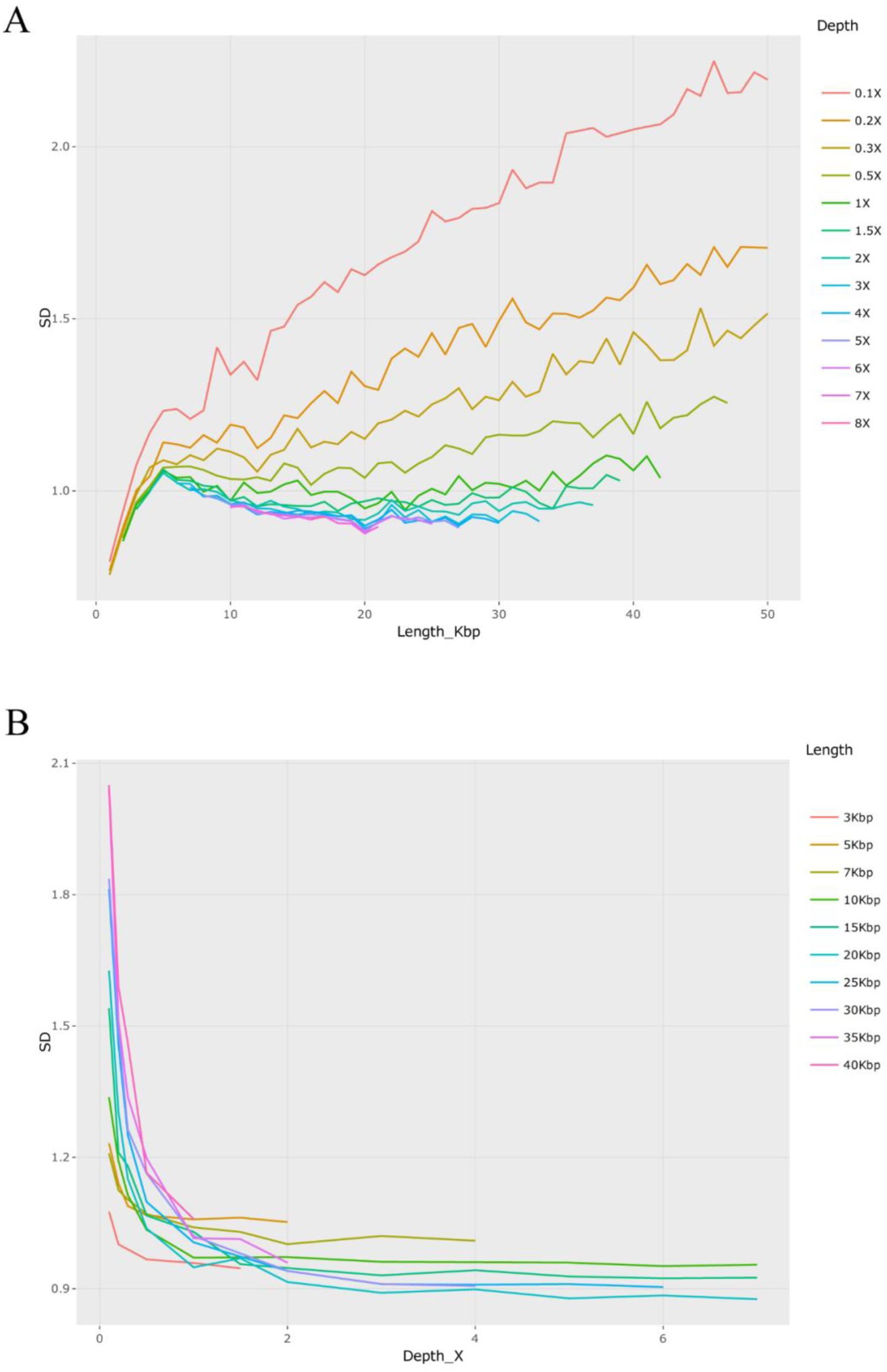
Ability to detect CNVs by low-coverage long-read sequencing data. To assess the effect of reads length and sequence depth on CNV calling, all the long reads from each individual were pooled together. Minimap2 was used for mapping all the long reads to human reference genome (GRCh37/hg19). We split the BAM file by lengths from 500bp to 30kb and randomly sample the split BAM file at different depth from 0.1X to 5X. Standard deviation (SD) of mean depth was calculated by a python scripts with 100kb window and 10kb sliding on genome sequence.

**Table S1.**
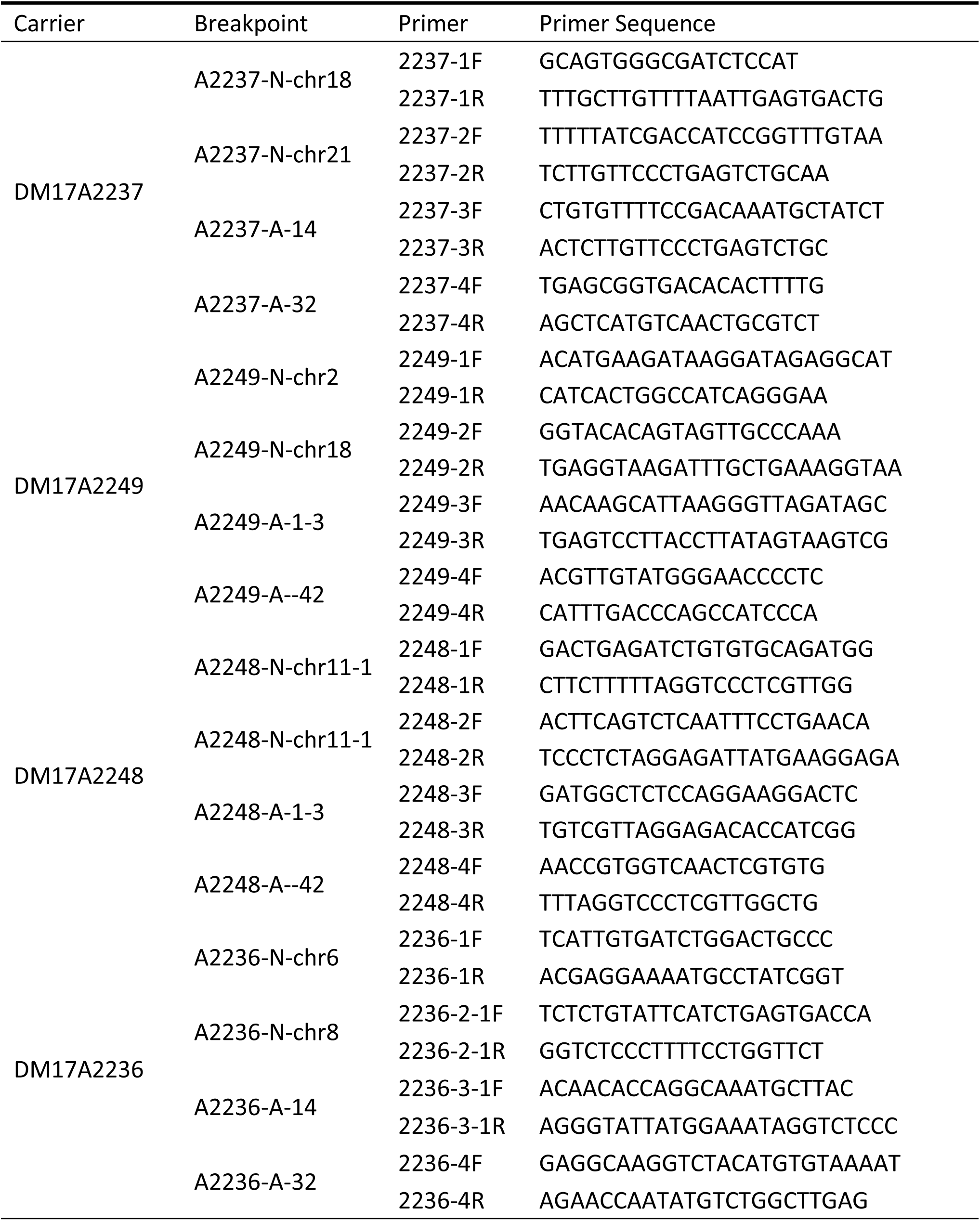
Design of PCR primers to validate translocations.

**Table S2.** Summary of long-read sequencing data on each subject.

| Sample | Cell number | Total Reads Bases | Total Reads Number | Pass Reads Bases | Pass Reads Number | Pass Reads Mean Length | depth(X) | Pass Reads N50 Length |
| --- | --- | --- | --- | --- | --- | --- | --- | --- |
| M17A2236 | 4 | 36,838,317,583 | 2,880,861 | 34,961,810,956 | 2,507,238 | 13,944 | 11.32 | 18,619 |
| DM17A2237 | 4 | 32,831,610,780 | 2,736,099 | 31,833,650,183 | 2,518,006 | 12,642 | 10.31 | 17,034 |
| DM17A2246 | 4 | 32,707,739,806 | 2,363,400 | 30,466,685,499 | 2,071,976 | 14,704 | 9.87 | 19,799 |
| DM17A2248 | 5 | 38,354,643,726 | 3,092,271 | 33,785,878,087 | 2,584,869 | 13,071 | 10.94 | 17,856 |
| DM17A2249 | 7 | 36,003,622,000 | 2,416,659 | 31,681,690,022 | 1,940,824 | 16,324 | 10.26 | 20,533 |
| DM17A2250 | 5 | 44,534,108,226 | 3,840,869 | 41,824,040,029 | 3,379,943 | 12,374 | 13.54 | 15,378 |
| DM17A2247 | 5 | 33,722,986,308 | 2,485,317 | 30,809,874,270 | 2,145,287 | 14,362 | 9.98 | 19,481 |

**Table S3.** Translocation detection and Breakpoint characterization by NGMLR and LAST in DM17A2246 and DM17A2247.

| Sample | Software | Mapping reads | Mapping ratio | Mapping reads | Mapping ratio | Mapping to different chromosome | Mapping to different chromosome |
| --- | --- | --- | --- | --- | --- | --- | --- |
| DM17A2246 | NGMLR | 1,931,841 | 93.24% | 129,684 | 6.71% | 36,560 | 1.89% |
|  | LAST | 1,988,419 | 95.97% | 762,114 | 38.33% | 47,821 | 2.40% |
| DM17A2247 | NGMLR | 2,024,874 | 94.39% | 133,608 | 6.60% | 38,813 | 1.92% |
|  | LAST | 2,070,876 | 96.53% | 811,189 | 39.17% | 46,538 | 2.25% |

